# A Novel Multilayer Cultivation Strategy Improves Light Utilization and Fruit Quality in Plant Factories for Tomato Production

**DOI:** 10.1101/2025.05.02.651818

**Authors:** Hanaka Furuta, Yuchen Qu, Dan Ishizuka, Saneyuki Kawabata, Toshio Sano, Wataru Yamori

## Abstract

Plant factories using artificial lighting are a promising solution to food security and urban agricultural challenges. However, cultivation of fruit-bearing crops such as tomatoes remains limited due to their high light demands, long growth periods, and tall plant structure. In this study, we aimed to develop an efficient cultivation system for tomatoes in a multilayer plant factory. Mini-tomatoes were hydroponically cultivated using white LEDs in a five-tier shelf system under two different cultivation methods. The conventional I-shaped method involved vertical growth on the top tier with downward lighting, while the novel S-shaped method trained each plant horizontally across the 2nd to 4th tiers with lateral lighting on each level. The S-shaped method enabled even light distribution, resulting in consistent photosynthetic rates throughout the canopy. In contrast, the I-shaped method suffered from strong light attenuation in the lower tiers, leading to reduced photosynthetic efficiency in shaded parts. Although total yield did not differ significantly between the two methods, the S-shaped method promoted earlier fruit maturation and improved fruit quality, including higher sugar content. Compared with greenhouse cultivation, plant factory conditions ensured stable temperature and lighting, leading to compact plant morphology, shorter internodes, and higher SPAD values. Moreover, fruit quality was more consistent year-round, with higher lycopene and sugar contents. This study demonstrates that the S-shaped cultivation system offers significant advantages in light use efficiency, plant management, and fruit quality. It represents a scalable approach for enhancing tomato production in plant factories and may facilitate the introduction of other high-light-demanding fruit crops into vertical farming systems.

## Introduction

According to a prediction by Benke and Tomkins (2018), the global population is expected to reach 9.7 billion. At the same time, increasing shortages of food and water, along with the loss of arable land due to climate change, continue to pose serious threats to human society, underscoring the urgent need to enhance agricultural productivity (Benke and Tomkins, 2018; Al-Kodmany, 2018; Kwon et al., 2020). Simultaneously, consumer demand for safer and healthier agricultural products is steadily growing (Ali et al., 2021). In response, recent advancements in plant factory technologies have gained significant attention and are emerging as a key component of sustainable agriculture.

Plant factories can be broadly classified into two types. Sunlight-type plant factories primarily utilize natural sunlight, with supplemental lighting provided under low-light conditions. In contrast, artificial light plant factories (ALPFs) rely entirely on artificial lighting sources. Both types use climate control systems (heating and cooling) and hydroponic techniques to ensure stable, year-round production and to minimize the risk of pests and contamination. Their enclosed environments allow for precise control of growth conditions, including light intensity, humidity, CO□ concentration, and nutrient delivery. As a result, plant factories are not affected by external weather or seasonal fluctuations and can achieve consistent yields (Takatsuji, 2010). Furthermore, their vertical space efficiency enables deployment in urban settings such as rooftops and indoor spaces where conventional farmland is unavailable (Al-Kodmany, 2018; Benke and Tomkins, 2018; SharathKumar et al., 2020). Controlled environments also allow for the optimized cultivation of high-value crops to meet consumer needs (Lee et al., 2014; Kang et al., 2016; Olvera-Gonzalez et al., 2021).

However, plant factories face two major challenges. First, they require high and continuous operational costs for lighting, temperature control, and water circulation. In Japan, although measures such as replacing fluorescent lights with LEDs and shifting lighting periods to nighttime (to take advantage of lower electricity rates) have been implemented, only 50–60% of facilities reportedly generate a positive net profit (Nousui, 2022). Second, plant factories are primarily designed for leafy vegetables, which have relatively low light requirements. The cultivation of fruit-bearing crops remains rare in ALPFs due to their longer growth periods, greater energy demands, and larger spatial requirements (Yoshita rt al., 2013). Nevertheless, recent efforts have explored the feasibility of producing strawberries, blueberries, and figs in plant factories (Yoshida et al., 2013; Aung et al., 2014; Kawamata et al., 2002). This emerging trend toward fruit crop production in plant factories holds great promise for improving food security and achieving sustainable production.

Tomatoes are among the most widely cultivated and consumed vegetables globally, with annual production exceeding 180 million tons according to the FAO (FAO STAT, 2020). In addition to fresh consumption, tomatoes are processed into sauces, ketchups, soups, and juices. They are also rich in bioactive compounds such as lycopene and GABA, which have been linked to health benefits (Palozza et al., 2011; Yamakoshi et al., 2007; Ali et al., 2021). However, tomato production is highly sensitive to environmental fluctuations. Temperatures below 14□°C or above 26□°C can inhibit stem, leaf, and flower development, reducing both yield and fruit quality (Adams et al., 2001; Shamshiri et al., 2018). Inconsistent or insufficient lighting can further degrade fruit quality, promote excessive vegetative growth, and reduce overall yield (Gomez and Mitchell, 2016; Paucek et al., 2020; López-Díaz et al., 2020; Wang et al., 2020). These issues can be addressed by growing tomatoes in plant factories, where environmental parameters can be tightly controlled.

Currently, tomato cultivation accounts for approximately 65% of production in sunlight-type plant factories. These facilities achieve stable yields even in regions with limited sunlight, such as the UK and Canada, by supplementing with artificial light (Liu et al., 2019). Numerous studies have reported that the use of LEDs as supplemental lighting enhances tomato growth, yield, and fruit quality (Adams et al., 2001; Gomez and Mitchell, 2016; Tewolde et al., 2018; Paucek et al., 2020; López-Díaz et al., 2020; Paponov et al., 2020). Nonetheless, tomato cultivation remains uncommon in ALPFs due to the crop’s high light demand and large spatial footprint. While dwarf tomato varieties have been proposed as a solution for vertical space efficiency and increased yield per area (Kwon et al., 2020; Kobayashi and Tabuchi, 2022; Ke et al., 2022), studies have shown that such varieties often produce fruit of lower quality compared to standard cultivars (Trien et al., 2021; Anu et al., 2021). Therefore, exploring viable cultivation methods for conventional tomato cultivars in ALPFs remains an important research priority.

In this study, we developed a new multi-tier cultivation system for tomato production in an ALPF. Unlike conventional vertical string trellising (I-shaped cultivation), our method employs an S-shaped growth strategy, in which tomato plants are guided through a curved, back-and-forth pattern across multiple layers of LED panels. This structured growth architecture enhances light interception, improves space efficiency, and contributes to improved yield and fruit quality in a closed, artificially lit environment.

## Results

### Environmental parameters of different cultivation methods

Temperature and light intensity were continuously monitored in the greenhouse. Both parameters exhibited rapid fluctuations across different months and throughout the day (Fig. 1A). In contrast, within the plant factory, temperature, light intensity, and air humidity were recorded continuously during the cultivation period at various plant levels. The average temperature in the plant factory showed minimal variation both seasonally and diurnally (Fig. 1B), and no significant differences in temperature or humidity were detected among the different plant levels in either the I-shape or S-shape cultivation systems (Fig. 1B). However, significant differences in light intensity were observed among the upper, middle, and lower plant levels under I-shape cultivation, while such differences were not evident in the S-shape system (Fig. 1B).

**Figure 1.**
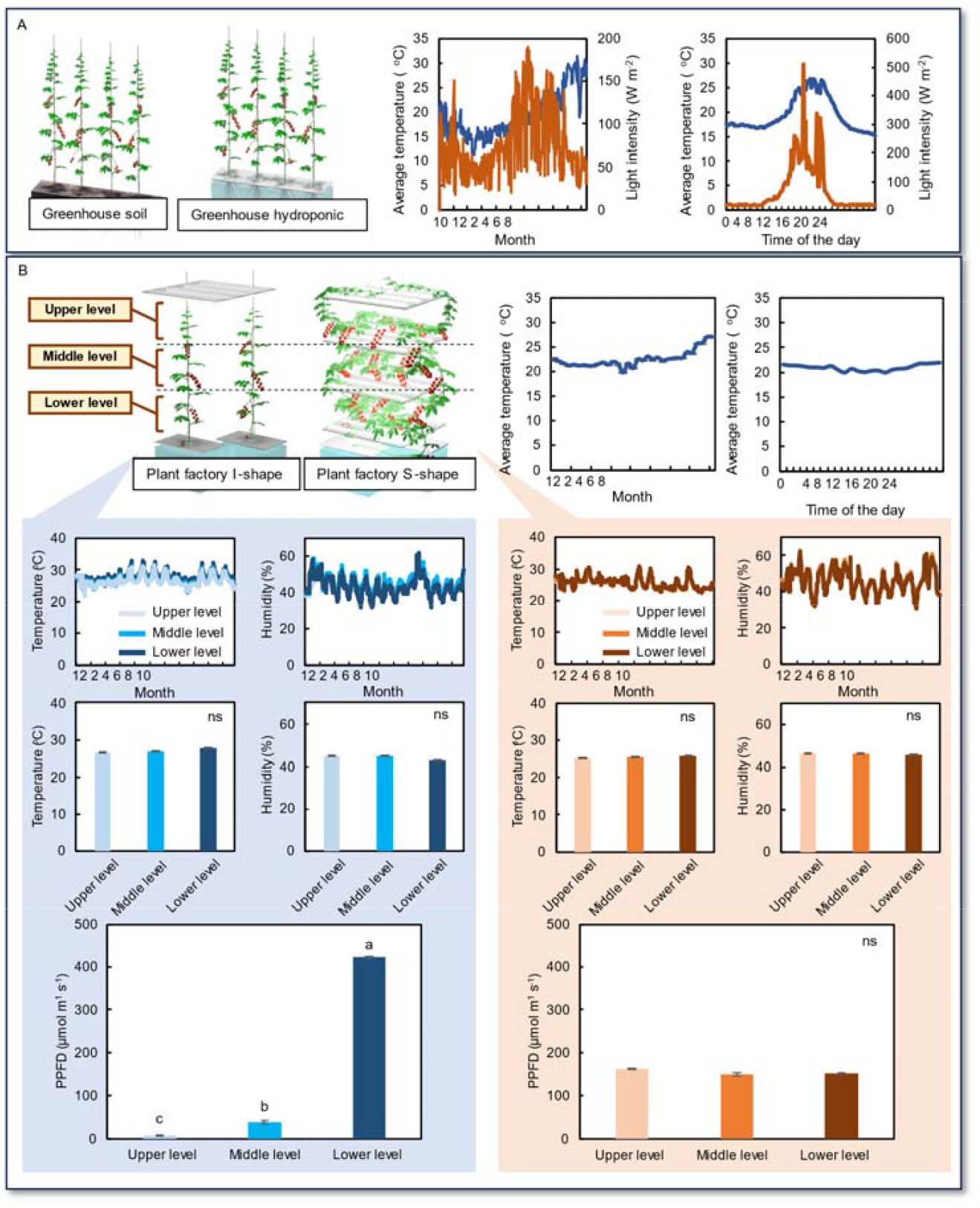
Environmental conditions in greenhouse and plant factory cultivation systems. **A**. Average temperature and light intensity in the greenhouse across the entire cultivation period and on a daily basis. **B**. Average temperature in the plant factory across the entire cultivation period and on a daily basis. In the plant factory, environmental parameters—temperature, humidity, and light intensity—were recorded separately at the upper, middle, and lower levels of the cultivation racks. Measurements were conducted for both the I-shape cultivation method, in which plants were trained vertically, and the S-shape cultivation method, in which plants were trained horizontally along multiple shelf levels to maximize spatial and light-use efficiency. Data is mean ± SE. Bars with the same letter are not significantly different n = 8.

### Photosynthetic and growth parameters of plants in different cultivation conditions

Photosynthetic parameters, including electron transport rate (ETR), photochemical quenching (1–qP), and non-photochemical quenching (NPQ), were determined at mature leaves located at the upper, middle, and lower levels across the various cultivation methods.

For greenhouse soil cultivation, clear differences in ETR were observed among the leaf levels: upper leaves exhibited the highest ETR, while lower leaves had the lowest (Fig. 2A). In contrast, the 1–qP parameter showed a reverse trend, with lower leaves displaying the highest values and upper leaves the lowest (Fig. 2A). However, NPQ did not differ significantly between the various leaf levels (Fig. 2A). In greenhouse hydroponic cultivation, the ETR values were similar between the upper and middle leaves, while the lower leaves showed a notably lower ETR (Fig. 2A). Similarly, the 1–qP values were higher in the lower leaves, whereas the middle and upper leaves exhibited comparable values (Fig. 2A). NPQ remained statistically consistent across all leaf levels in this cultivation method (Fig. 2A).

**Figure 2.**
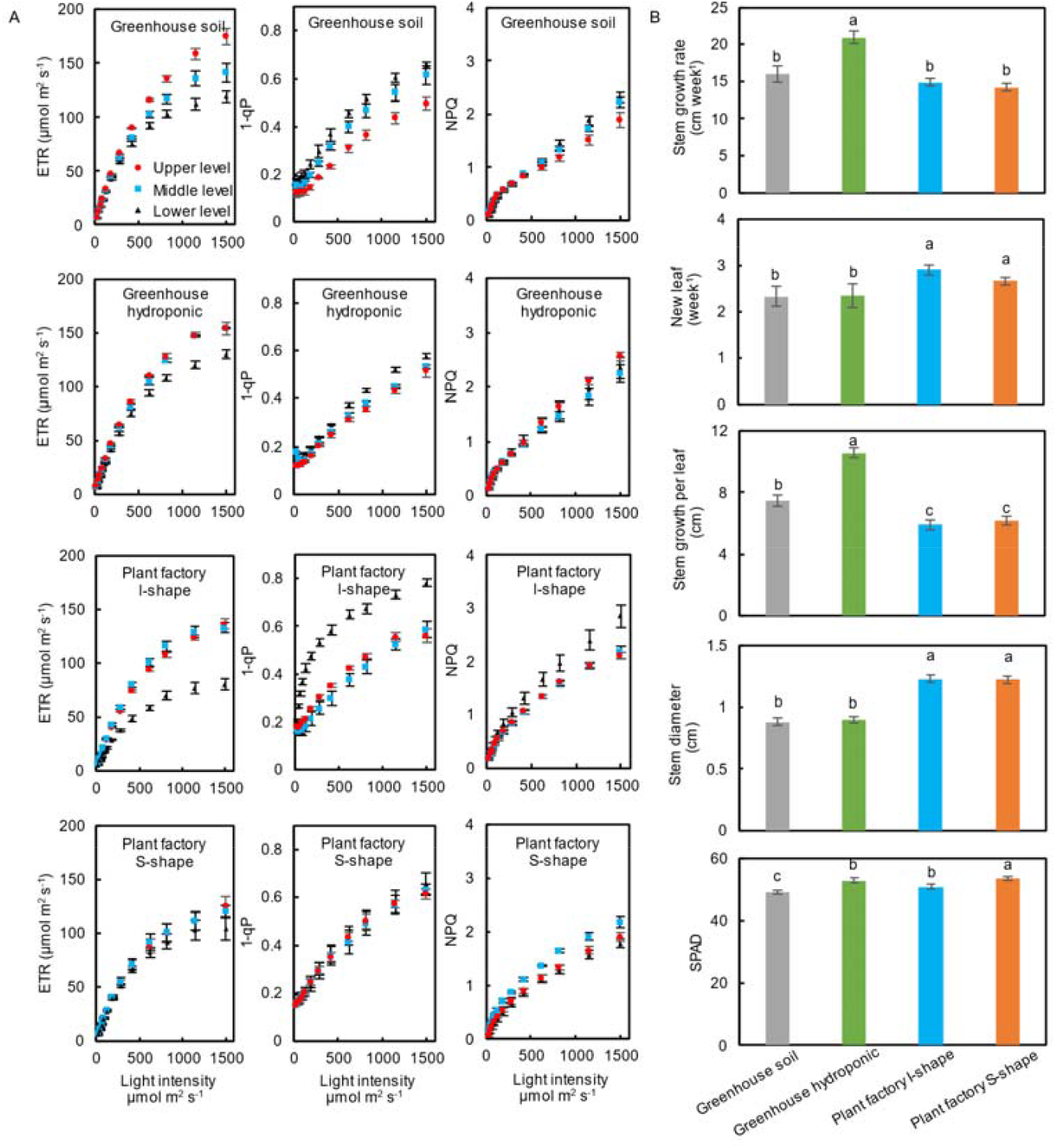
Photosynthetic and growth parameters of tomato plants under different cultivation conditions. **A**. Electron transport rate (ETR), photochemical quenching (1–qP), and non-photochemical quenching (NPQ) of tomato plants grown under four cultivation systems: greenhouse soil, greenhouse hydroponics, plant factory with I-shape cultivation, and plant factory with S-shape cultivation. **B**. Stem elongation rate, number of newly emerged leaves, Soil Plant Analysis Development (SPAD) value, and stem diameter of tomato plants cultivated under the same four conditions. Data is mean ± SE. Bars with the same letter are not significantly different n = 8.

In the plant factory under I-shape cultivation, the results mirrored those of the greenhouse hydroponic method: the lower-level leaves had a reduced ETR and elevated 1–qP in comparison to the upper and middle leaves, which maintained similar values (Fig. 2A). Again, NPQ did not show any significant differences among leaf levels (Fig. 2A). Conversely, under plant factory S-shape cultivation, no substantial differences in ETR or 1–qP were observed among the upper, middle, and lower leaves, although the middle leaves exhibited a slightly higher NPQ compared with the other levels (Fig. 2A).

When comparing overall plant growth characteristics among the cultivation methods (Fig. 2B), greenhouse hydroponic systems showed the highest stem growth rate. In contrast, the plant factory cultivations (both I-shape and S-shape) presented with a higher rate of new leaf emergence and a greater stem diameter than their greenhouse soil and hydroponic counterparts. Consequently, the average distance between leaves was greatest in the greenhouse hydroponic setup, whereas both plant factory systems had significantly lower inter-leaf distances. Additionally, the plant factory S-shape cultivation yielded the highest SPAD values, while the greenhouse soil cultivation displayed the lowest (Fig. 2B).

### Productivity and fruit quality parameters of plants in different cultivation conditions

Fruit yield data were collected throughout the cultivation period to assess both productivity and fruit quality across all methods. Although no significant differences were found in yield per cultivation area, per plant, or per cluster among the different methods (Fig. 3A), both plant factory S-shape and I-shape systems demonstrated a significantly shorter harvest cycle, from the first to the 23rd cluster, compared with the greenhouse cultivations. Among these, the plant factory S-shape system exhibited the shortest harvest cycle (Fig. 3A).

**Figure 3.**
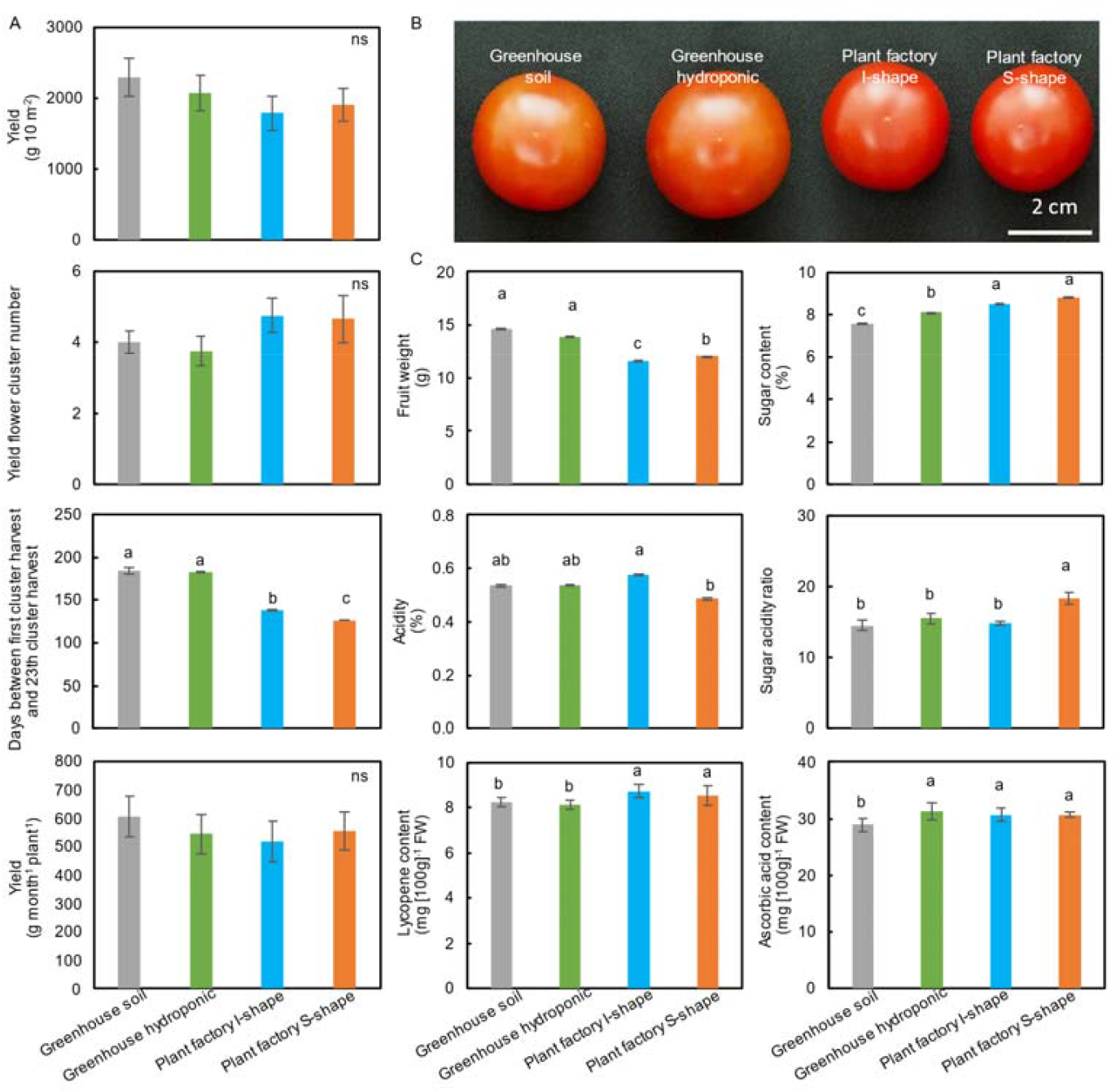
Productivity and fruit quality of tomato plants cultivated under different conditions. **A**. Productivity parameters including yield per unit area, yield per flower cluster, yield rate, and monthly yield per plant across different cultivation systems. **B**. Representative images of tomato fruits harvested from each cultivation method. **C**. Fruit quality parameters including average fruit weight, acidity, soluble sugar content, lycopene concentration, and ascorbic acid content under each cultivation condition. Data is mean ± SE. Bars with the same letter are not significantly different n = 50 - 90.

In terms of fruit appearance and weight, fruits from plant factory I-shape and S-shape systems were smaller and had a deeper red color compared with those from the greenhouse (Fig. 3B). Further analysis revealed that fruits harvested from the plant factory systems had significantly lower fruit weights than those from the greenhouse, with the I-shape system showing the lowest fruit weight overall (Fig. 3B).

Regarding fruit quality, plant factory tomatoes exhibited significantly higher sugar content compared with greenhouse tomatoes, with the greenhouse soil method yielding the lowest sugar levels (Fig. 3C). Additionally, the plant factory S-shape system produced tomatoes with the lowest fruit acidity, resulting in the highest sugar–acidity ratio, while the other methods showed higher acidity and lower sugar–acidity ratios (Fig. 3C). Finally, plant factory tomatoes had significantly higher lycopene content than those from the greenhouse, and ascorbic acid levels were similar across all treatments except for the greenhouse soil cultivation, which had a significantly lower value (Fig. 3C).

### Key-bioactive compounds within metabolism map

Eight amino acids in the fruit were quantified to assess the effects of different cultivation systems on nutrient composition. Overall, tomatoes grown in the greenhouse exhibited higher amino acid levels, particularly those directly linked to or derived from glycolysis. Within the greenhouse systems, soil cultivation resulted in higher concentrations of phenylalanine and serine, while the hydroponic system produced higher levels of aspartic acid, GABA, glutamic acid, and proline (Fig. 4). A notable exception was asparagine: both greenhouse soil and plant factory S-shape cultivations resulted in elevated asparagine content, whereas greenhouse hydroponic cultivation showed the lowest level among all treatments (Fig. 4).

**Figure 4.**
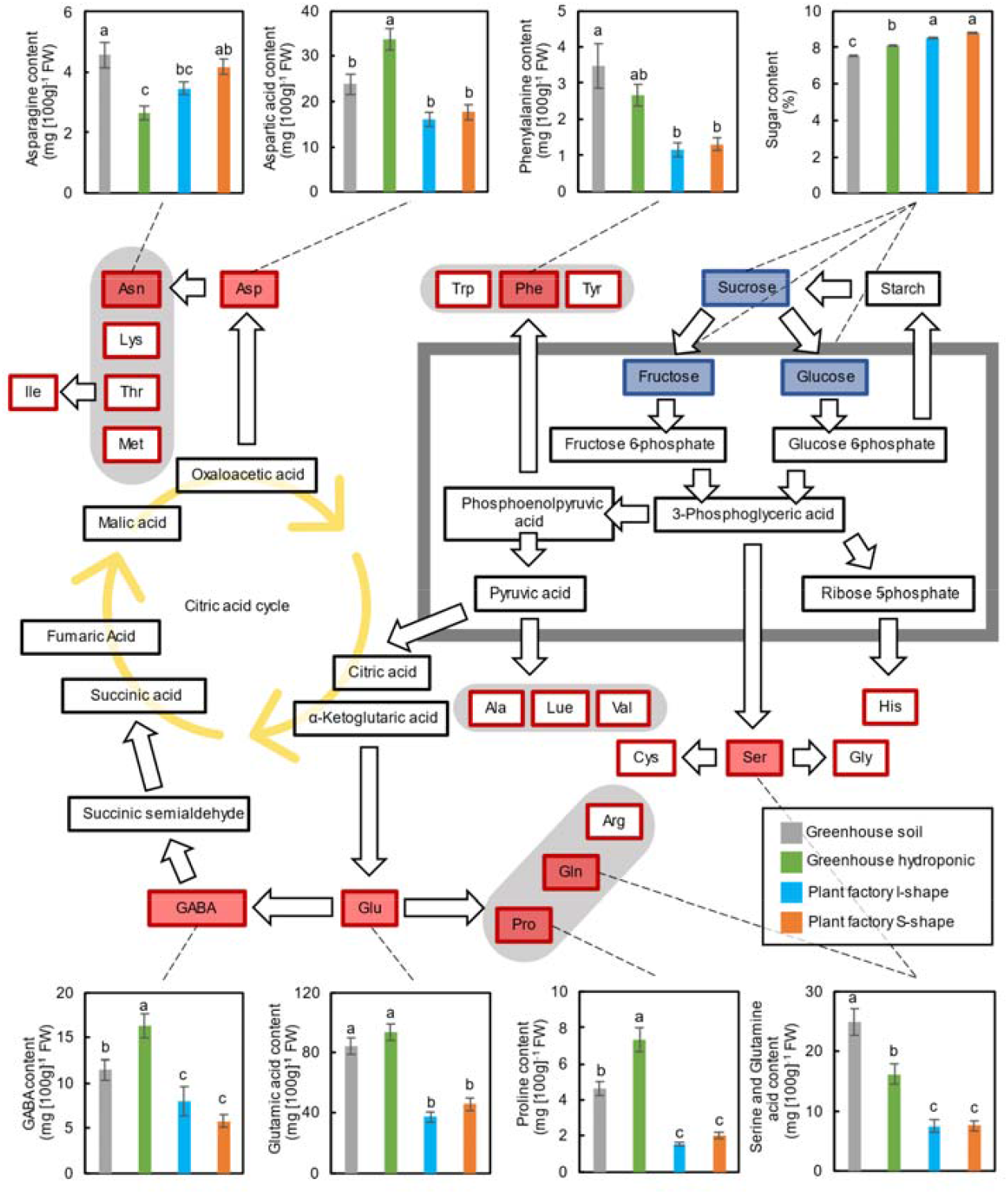
Metabolic map of amino acids and sugars in tomato. The contents of amino acids (indicated by red squares) and sugars (indicated by blue squares) were quantified under different cultivation conditions and are shown in the corresponding bar graphs. Data is mean ± SE. Bars with the same letter are not significantly different n = 50 - 90.

## Discussion

This study aimed to establish a novel tomato cultivation method optimized for closed-type plant factories with artificial lighting. Unlike previous approaches that rely on dwarf cultivars with limited quality (Trien et al. 2021; Anu et al. 2021), we developed a system utilizing high-quality commercial varieties commonly used in greenhouse or open-field cultivation. Two cultivation systems were tested: the conventional vertical “I-shape” method and a novel space-efficient “S-shape” system leveraging multi-layered cultivation shelves. Compared to greenhouse cultivation, both methods led to higher inflorescence numbers, increased sugar and lycopene content, with the S-shape system further enhancing sugar-acid balance. Generally speaking, plant factory cultivations showed higher growth and ripe speed, and higher sugar contents, while greenhouse cultivations showed higher animo acid content (Fig 5), which represent interesting trade-off options for people to choose from based on their needs. Our results demonstrate the potential of this cultivation method to improve both yield and fruit quality in plant factories, paving the way for broader application to other fruiting vegetables and even space agriculture.

**Figure 5.**
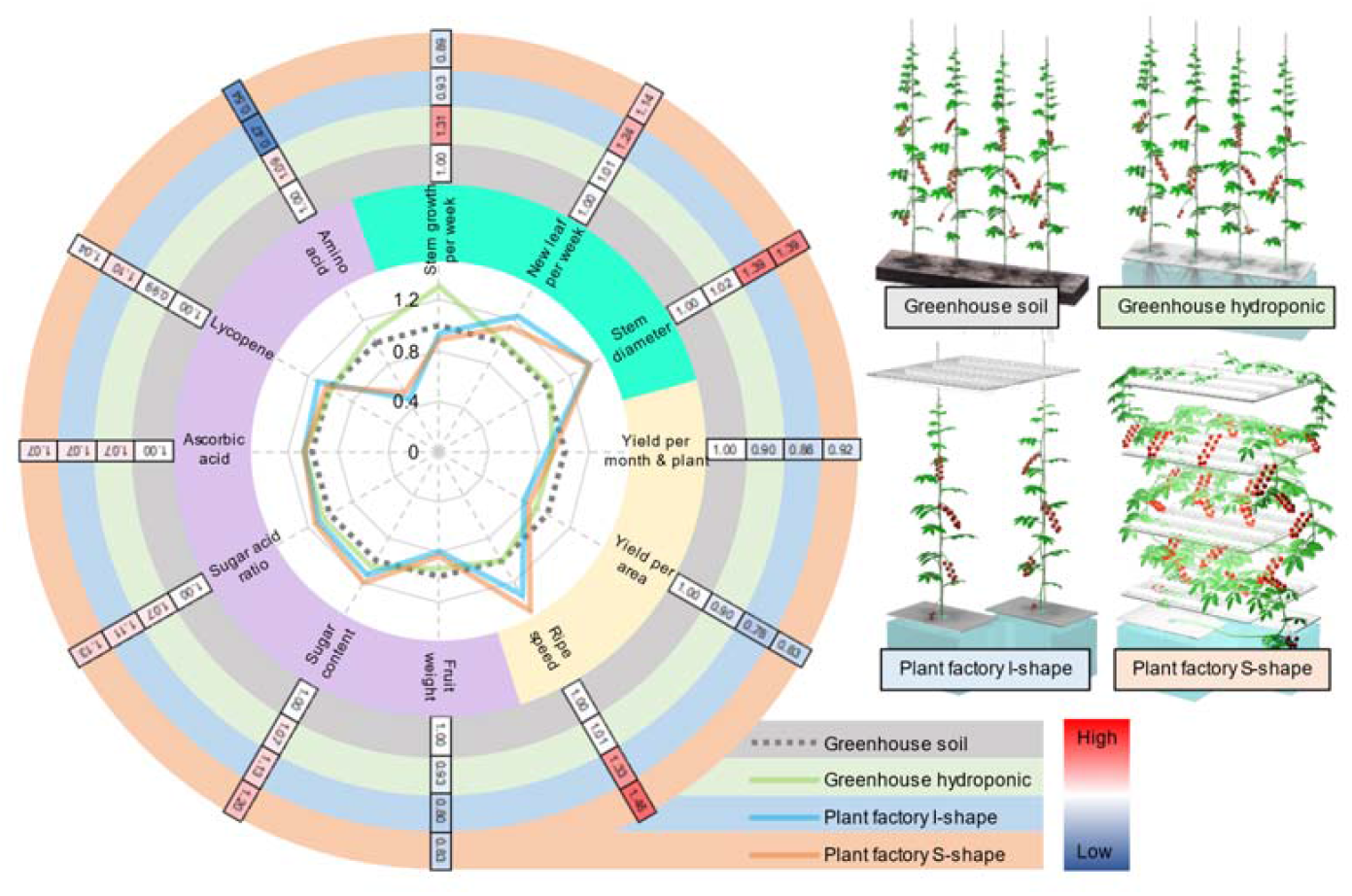
Comparison of key growth, productivity, and fruit quality traits of tomato plants under different cultivation conditions. The radar chart on the left displays relative values of growth parameters (mint green zone), productivity parameters (light yellow zone), and fruit quality parameters (plum zone), normalized against those under greenhouse soil cultivation. The outer heat map shows the actual values for each trait, with colored rings indicating the corresponding cultivation conditions.

### Enhanced Spatial Efficiency and Quality Stability of Tomato via “S-shape” system in Artificial Light-Type Plant Factories

In this study, we demonstrated that a novel S-shaped cultivation system for tomato cultivation in closed-type plant factories under artificial lighting improves plant growth and fruit quality compared to conventional I-shaped systems and greenhouse cultivation. Consistent with previous studies (Adams et al., 2001; Tewolde et al., 2018; Paucek et al., 2020; López-Díaz et al., 2020; Paponov et al., 2020), supplemental lighting in the middle and lower canopy enhanced light use efficiency, resulting in thicker stems, shorter internodes, higher chlorophyll content (SPAD values), and increased inflorescence number (Fig 2B, Fig S1). These morphological improvements led to stable fruit development and yield comparable to greenhouse systems, independent of season or weather conditions (Fig 3A, Fig S2).

High soluble solids content (°Brix) was observed particularly under the S-shaped system (Fig 3C, Fig S3, S4), likely due to enhanced photosynthetic activity across the whole plant (Fig 2A, Fig S1). This aligns with reports showing that increased light exposure improves sugar accumulation while reducing organic acid levels (Gomez and Mitchell, 2016; Gautier et al., 2009). Although no significant differences in ascorbic acid content were detected among treatments (Fig 3C, Fig S5), light exposure to the fruit surface—known to affect ascorbic acid biosynthesis (Gautier et al., 2009; Ntagkas et al., 2019)—could be better optimized in future trials by adjusting leaf arrangement or light positioning.

Lycopene content was higher in plant factory-grown tomatoes than in greenhouse-grown ones (Fig 3C, Fig S5A, B), possibly due to the stable light and temperature conditions mitigating the suppressive effects of high fruit surface temperature on lycopene biosynthesis (Helyes et al., 2007; Bianchetti et al., 2020) (Fig 1B, Fig S1). However, no significant difference was observed between I- and S-shaped systems (Fig 3C, Fig S5A, B). Future studies using controlled spectral light (e.g., red/blue LEDs) and fresh fruit measurements could further clarify these effects.

Interestingly, the concentrations of key amino acids such as glutamate and GABA were lower in plant factory-grown fruit (Fig 4, Fig S5C). One plausible explanation is reduced carbon flux into amino acid biosynthesis due to enhanced sugar retention under controlled lighting. Previous studies have shown that fruit amino acid profiles are sensitive to light intensity, spectral quality, and CO□ levels (Dhakal and Baek, 2014; Ntagkas et al., 2020; Yan et al., 2021), underscoring the need for detailed environmental monitoring around the fruit.

Overall, our findings suggest that the S-shaped cultivation method, by maximizing spatial and light use efficiency, not only improves yield and sugar content but also offers a promising strategy for stable, high-quality tomato production in vertical farming systems and potential applications in resource-limited environments such as space agriculture.

### Environmental and Physiological Factors Affecting Fruit Quality Under Artificial Lighting Conditions

Machine learning models using environmental variables—such as radiation, temperature, humidity, CO□ concentration, and leaf area—have demonstrated the ability to predict fruit quality attributes like soluble solids and glutamate content regardless of tomato cultivar or origin (Xiang et al., 2022; Yoshida et al., 2020). In this study, decision tree regression analysis incorporating plant growth parameters and environmental data revealed that fruit quality traits were primarily influenced by stem diameter and SPAD values (Fig S6). Specifically, soluble solids were correlated with stem diameter and SPAD (R^2^ = 0.34), ascorbic acid with SPAD and internode length (R^2^ = 0.31), and glutamate and aspartate with stem diameter (R^2^ = 0.64 and 0.51, respectively) (Fig S6).

These findings suggest that improved photosynthetic activity in leaves, facilitated by stable light conditions in plant factories, enhances sugar accumulation, as previously reported (Gomez and Mitchell, 2016; Paucek et al., 2020). Furthermore, fruit-localized light sensing through phytochrome has been shown to suppress starch synthesis and promote sugar accumulation in tomato fruits (Bianchetti et al., 2018), emphasizing the potential importance of direct fruit irradiation.

Our analysis also indicated that environmental variables such as cumulative temperature and radiation were strong predictors of plant growth (Fig S6), particularly SPAD values and stem thickness (R^2^ = 0.86). However, several fruit metabolites, such as organic acids and lycopene, were not well explained by plant growth traits alone. Previous studies have demonstrated that lycopene accumulation is enhanced by red and blue light (Xie et al., 2019; Zhang et al., 2020), and reduced by elevated fruit surface temperatures (Helyes et al., 2007), highlighting the dual influence of light quality and heat. To directly assess the effect of fruit irradiation, we conducted a light-shielding experiment using neutral density filters (Fig S7). While fruit surface temperature was only marginally affected (Fig S7A and B), both ascorbic acid concentration and lycopene content significantly increased in fruits exposed to higher light levels (Fig S7C). These results provide experimental evidence that light exposure to fruit tissue itself, not just to leaves, is a critical factor for promoting the biosynthesis of health-promoting compounds such as antioxidants and carotenoids.

In conclusion, integrating light distribution models that account for both leaf and fruit exposure may enhance the optimization of controlled-environment tomato production systems. Our findings underscore the potential of artificial light-based cultivation not only to stabilize yield but also to improve functional quality, which is particularly relevant for plant factories aiming for high-value crop production.

### Yield Maximization through Sequential Planting in Vertical S-shaped Cultivation

In this study, we demonstrated that the S-shaped cultivation system under artificial lighting conditions achieved tomato yields comparable to those obtained through conventional greenhouse cultivation (Fig. 3A). To further enhance productivity per unit area, we conducted a yield simulation incorporating staggered, sequential planting within the same vertical cultivation space. During cultivation, we observed that the lower stem regions beneath the uppermost harvested truss were mostly defoliated and underutilized. We hypothesized that this unoccupied space could be repurposed for additional growth by either cultivation axillary shoots or transplanting a second seedling, both following the same S-shaped pattern (Fig. S8). This approach aims to maximize spatial and energy efficiency while increasing total yield within a fixed cultivation period.

Based on plant growth patterns, we estimated that initiating the growth of an axillary shoot or introducing a secondary seedling approximately 10 weeks after the initial planting would align the shoot apex of the second plant with the height of the uppermost harvested truss of the first plant. At that stage, aged leaves of the primary plant could be pruned following fruit harvest to free up space and enhance light penetration to the developing apex of the second plant (Fig. S8). This strategy would enable both plants to be trained within the same frame without spatial interference. Simulations based on empirical cultivation data predicted that the introduction of a second plant could increase the total yield per cultivation unit from 2.76 kg to 4.05 kg, representing a 1.47-fold increase in yield per square meter, from 9.50 to 13.94 kg m□^2^.

These findings highlight the potential of time-staggered interplanting as a strategy to maximize spatial efficiency in vertical plant factory systems. Further experimental validation is needed to assess the physiological impact of overlapping root and shoot systems, including possible competition for light and nutrients. Moreover, scalability using larger rack systems and optimization of planting intervals could further enhance the productivity of this cultivation method.

### Potential Application of the S-shaped Cultivation Method in Space Agriculture

Controlled environment agriculture using artificial light has been increasingly recognized as a promising solution for future space missions. In long-duration spaceflights, pre-packaged meals often degrade in quality and lack sufficient essential nutrients, which may compromise astronaut health and appetite due to so-called “menu fatigue” (Khodadad et al., 2020; Sirmons et al., 2020). To address this, NASA’s “VEGGIE” project has conducted several successful experiments growing leafy greens and dwarf tomato cultivars on the International Space Station (NASA, 2022). Our study demonstrates a novel S-shaped tomato cultivation method using standard high-quality varieties rather than dwarf types. This system efficiently utilizes vertical space and artificial light (Fig 1B), enabling stable fruit quality and yield under completely controlled conditions. Importantly, our method allows for precise regulation of environmental factors such as light spectrum and intensity, which can be leveraged to optimize growth rates and enhance accumulation of beneficial compounds such as lycopene and ascorbic acid—key nutrients often limited in space diets.

Given these advantages, the S-shaped cultivation method developed here could serve as a model for high-efficiency crop production systems in extraterrestrial habitats. Furthermore, its applicability to other fruiting crops offers a foundation for developing integrated plant-based life support systems capable of providing a balanced diet for astronauts during extended missions.

## Material and Method

### Plant material and cultivation

In this study, the mini tomato variety CF Chika (*Solanum lycopersicum*), developed by Takii Seed Company, was used. For soil-based greenhouse cultivation, a base fertilizer application was conducted using 1 kg of nitrogen-phosphorus-potassium (NPK) mixed fertilizer (N=12, P=8, K=10) and 1 kg of carbonate magnesium lime per 10 square meters (1m × 10m) of cultivation area. For hydroponic cultivation in greenhouse, plants were grown in a substrate of crushed coconut shell, which serves as a sustainable and well-aerated growing medium. The coconut shell is enclosed within hydrophilic textiles that partially contact a nutrient solution reservoir below. This setup allows water and nutrients to be drawn up through capillary action, ensuring a consistent supply to the plant roots. Additionally, two drip irrigation tubes are embedded within the coconut shell medium to provide supplementary nutrient delivery, optimizing moisture distribution. Any excess nutrient solution drains back into the reservoir, where it is collected and recirculated. For hydroponic cultivation in plant factory, plants are cultivated with their roots fully submerged in a continuously circulated nutrient solution tank. All nutrient solution mentioned above were made with commercial liquid fertilizer with balance NPK, and maintained at an electrical conductivity (EC) of 1.5 ± 0.05 dS m^-1^. Nutrient solutions were regularly replaced to remove metabolic waste produced by plants.

For both greenhouse and plant factory, environmental temperature and illumination were recorded every 1 hour with TR-76Ui - USB Connectible CO2 Logger (T&D Corporation) and e-kakashi (PS Solutions Corporation – Agricultural IoT Solutions). In order to study the relationship between illumination and fruit surface temperature, some fruit clusters were covered with shading filters with transparent rates of 69.3% and 6.6% (LEE FILTER). Plant surface temperature was determined with a thermography camera (FLIR C3).

### Set up of I shape and S shape cultivation system

Tomato plants in greenhouse were string trellised to grow vertically (I-shape). Tomato plants in plant factory was cultivated within a 135 cm x 67 cm x 200 cm frame which was used to install illumination devices and string trellising. For I shape cultivation, tomato plants were guided to grow vertically within the frame and illumination device was installed at the very top of the frame (Fig 1A). For S shape cultivation, tomato plants were guided to grow back and force through multiple layers within the frame, and illumination devices were installed at each layer (Fig 1B).

Both I shape and S shape cultivations in plant factory were illuminated with white light LED at a 16-hour/8-hour light-dark circle. For I shape cultivation, light intensities 35 cm, 70 cm and 105 cm below illumination devices were determined to be 422 μmol m^-2^ s^-1^, 38 μmol m^-2^ s^-1^ and 7 μmol m^-2^ s^-1^ during cultivation with the present of tomato plants. For S shape cultivation, illumination devices were installed at every layer between stem of tomato plants, result in a light-plant-light-plant multilayer sandwich structure (Fig 1B). Light intensities in the middle area between each two layers of illumination devices were determined to be 150 ±3 μmol m^-2^ s^-1^ uniformly, with the present of tomato plants.

### Determination of photosynthetic parameters

Real-time photosynthetic fluorenes in PSII were determined with MICRO-PAM device (Walz) in cultivation conditions. Measurements were taken at higher section (30 cm below shoot apex), middle section (60 cm below shoot apex) and lower section (90 cm below shoot apex). The effective quantum yield of photochemical energy conversion during PSII (Y[II]), the non-photochemical quenching (NPQ), the photochemical quenching of PSII (qP) was calculated as follows according to Alexander V. Ruban (2017). The electron transport rate (ETR) of PSII and PSI were calculated as ETR I (or ETR II) =□0.5□×□abs I ×□Y(I) (or Y[II]), where 0.5 is the fraction of absorbed light between PSII and PSI (assuming they are equal), and abs I is absorbed irradiance taken as 0.84 of incident irradiance. ETR were also determined throughout light period in certain days during cultivation, and the data was recorder every minutes. The content of chlorophyll was also determined with a hand-held SPAD-502 meter (Spectrum Technologies, Inc.) at higher section, middle section and lower section.

### Determination of plant growth and fruit quality

Tomato plants stem length, stem diameter, fully expanded (main axis longer than 10 cm) leaf number, leaf length and SPAD value of the leave right above all flower clusters were recorded every week to determine plant growth.

Ripened fruits were harvested for the determination of fruit quality. Mini tomato usually yields 10 fruits from 1 cluster, but the very fruit adjacent to stem frequently suffer from malformation. Therefore only 9 fruits from 1 cluster were used for fresh weight determination. 3 fruits from every cluster were chosen and cut into half. Half of the fruits were used for the determination of sweetness (Brix) and acidity determination with ATAGO PAL-BX|ACID3 Pocket Brix-Acidity Meter (ATAGO Co., Ltd.) and determination of L-ascorbic acid with test paper (Reflectoquant, Merck KGaA). Lycopene was determined as described by Ito and Horie, 2009. The rest half of the fruits were used for the quantification of amino acid including Aspartate, Glutamate, Asparagine, Serine, Glutamine, Proline, Gamma-Aminobutyric Acid (GABA) and Phenylalanine through pre-column derivatization using phenyl isothiocyanate (PITC), followed by separation and detection via High-Performance Liquid Chromatography (HPLC), as described by Heinrikson and Meredith (1984).

### Statistical analyses

Data for growth determination including stem length, stem diameter, leaf number, leaf length and SPAD value were collected throughout 48 weeks of cultivation in greenhouse, and 30 weeks of cultivation in plant factory. Data for fruit fresh weight and yield determination were collected from 35 clusters in greenhouse soil cultivation, 36 clusters in greenhouse hydroponic cultivation, and 23 clusters in plant factory cultivation.

## Acknowledgments and Funding

This work was supported by KAKENHI (18KK0170, 21H02171, and 24H02277 to W.Y.) from the Japan Society for the Promotion of Science (JSPS).

## Contributions

H.F. and W.Y. designed the experiments. H.F., Y.Q., D.I. and S.K. grew the plants, performed the experiments, and analyzed the data. T.S. performed quantifications of free amino acids and isoflavones. Y.Q. and W.Y. prepared the manuscript. All the authors have read and approved the final version of this manuscript. Author H.F. and Author Y.Q. contributed equally to this work and share first authorship.

## Data availability statement

Supporting data can be requested by contacting the corresponding author.

## Conflict of interests

The authors declare no conflict of interest.

## Notes

### Competing Interest Statement

The authors have declared no competing interest.

