## Supplementary Figure 1-8 for "A Novel Multilayer Cultivation Strategy Improves Light Utilization and Fruit Quality in Plant Factories for Tomato Production"

This document contains following materials:

Supplementary Figure 1-8

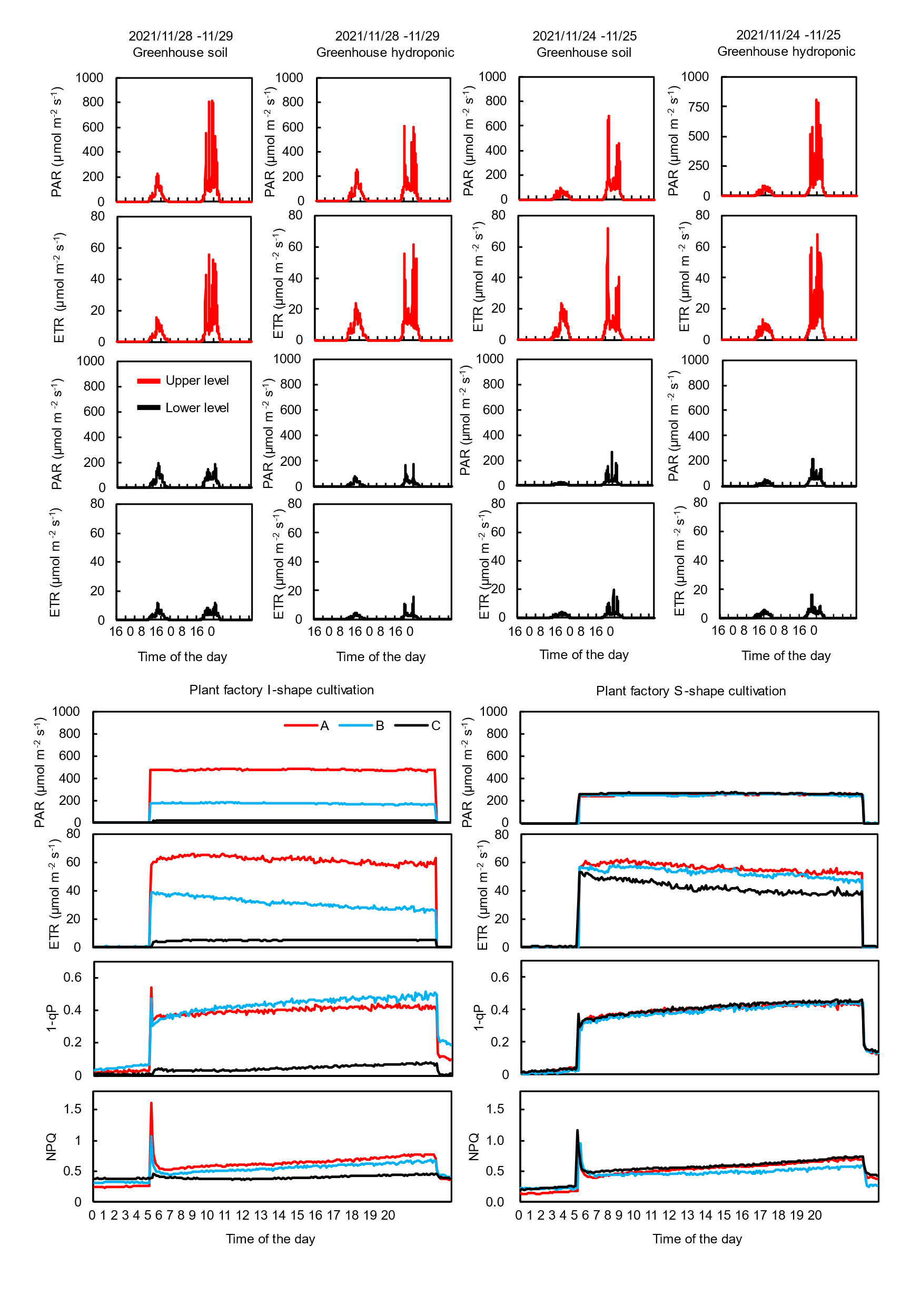

Figure S1

Record of light intensity and corresponding photosynthetic activities at different levels of plant leaves in greenhouse and plant factory cultivation throughout two days.

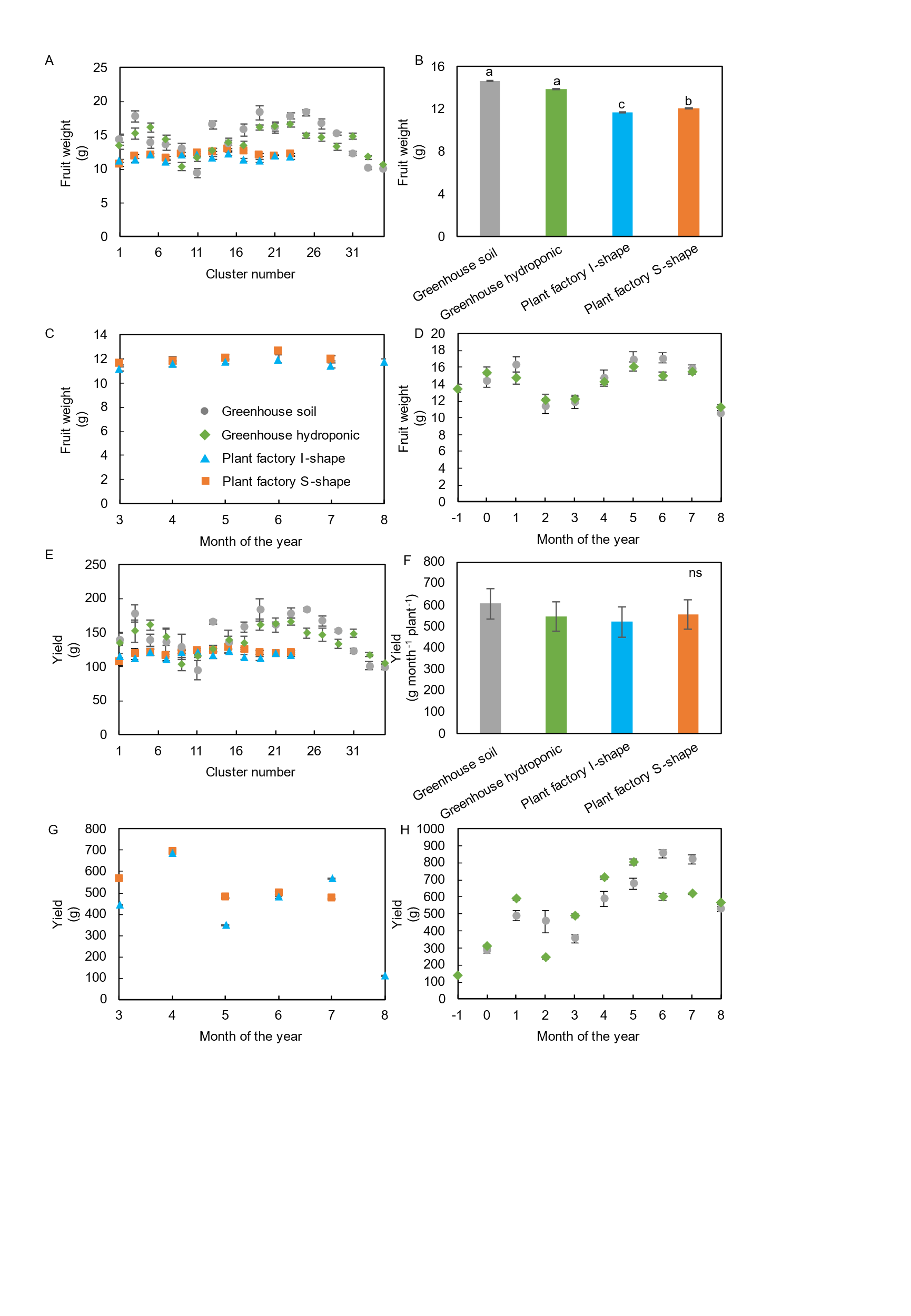

Figure S2

Fruit weight and yield weight of tomato plants cultivated in different conditions. A. Fruit weight of each cluster. B. Average fruit weight. C, D. Fruit weight at different month of a year. E. Yield of each cluster. F. Average yield. G, H. Yield at different month of a year. Data is mean ± SE. Bars with the same letter are not significantly different n = 8.

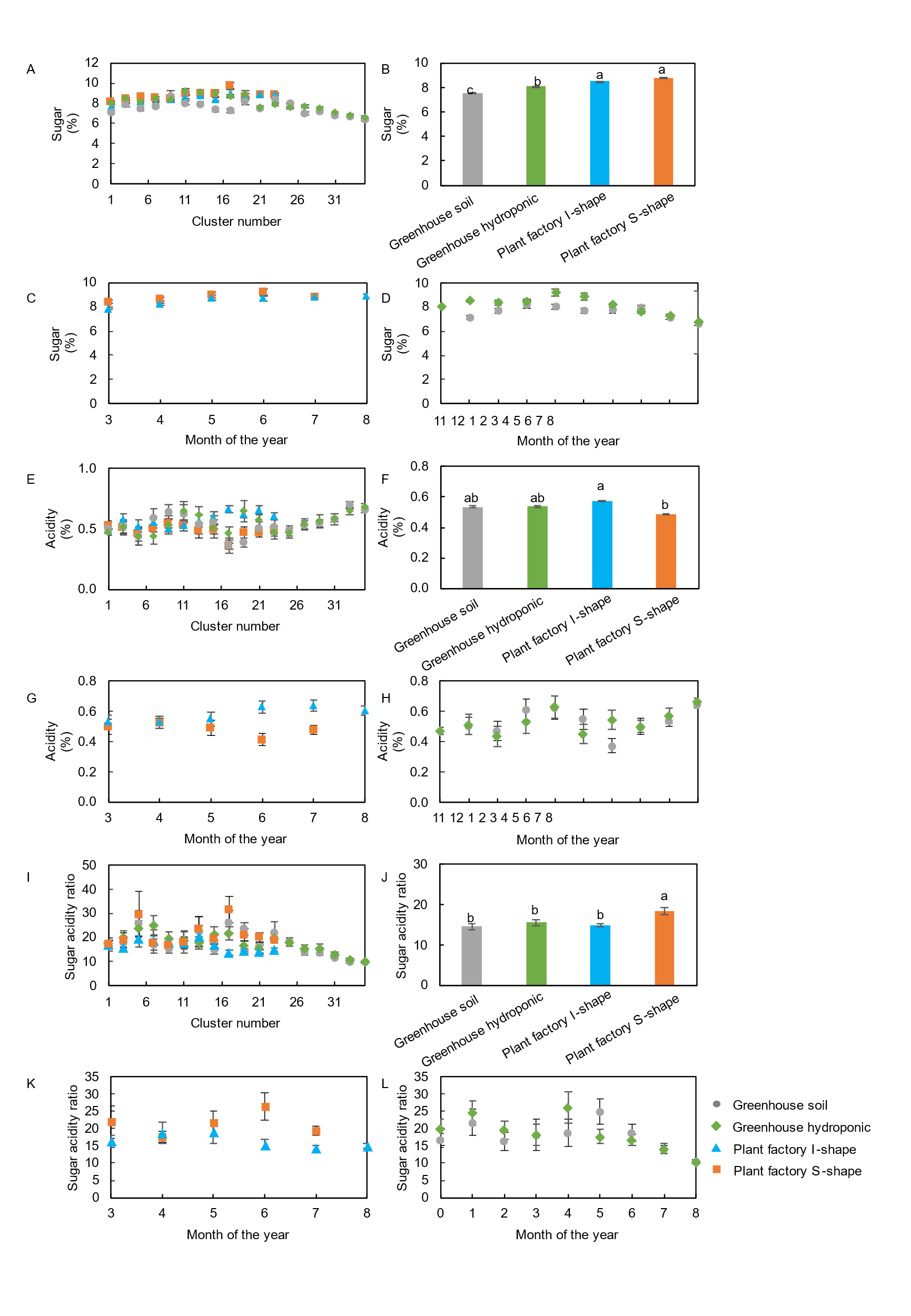

Figure S3

Sugar and acidity of tomato plants cultivated in different conditions. A. sugar content of each cluster. B. Average sugar content. C, D. Sugar content at different month of a year. E. Acidity of each cluster. F. Average acidity. G, H. Acidity at different month of a year. I-L. Sugar/acidity ratio tomato fruits. Data is mean ± SE. Bars with the same letter are not significantly different n = 8.

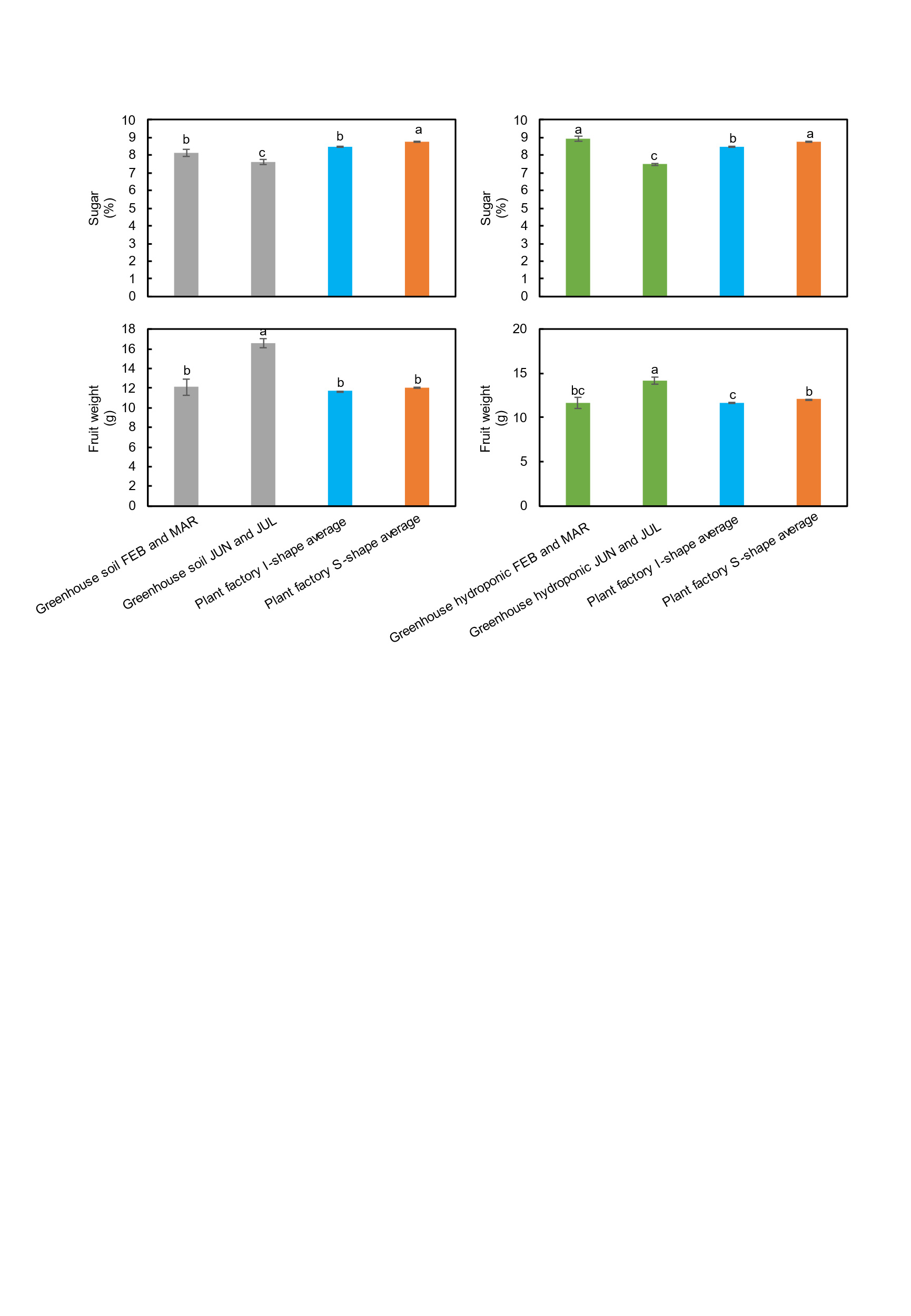

Figure S4

Sugar content and fruit weight of greenhouse between different months of a year compared with plant factory. Data is mean ± SE. Bars with the same letter are not significantly different n = 50-90.

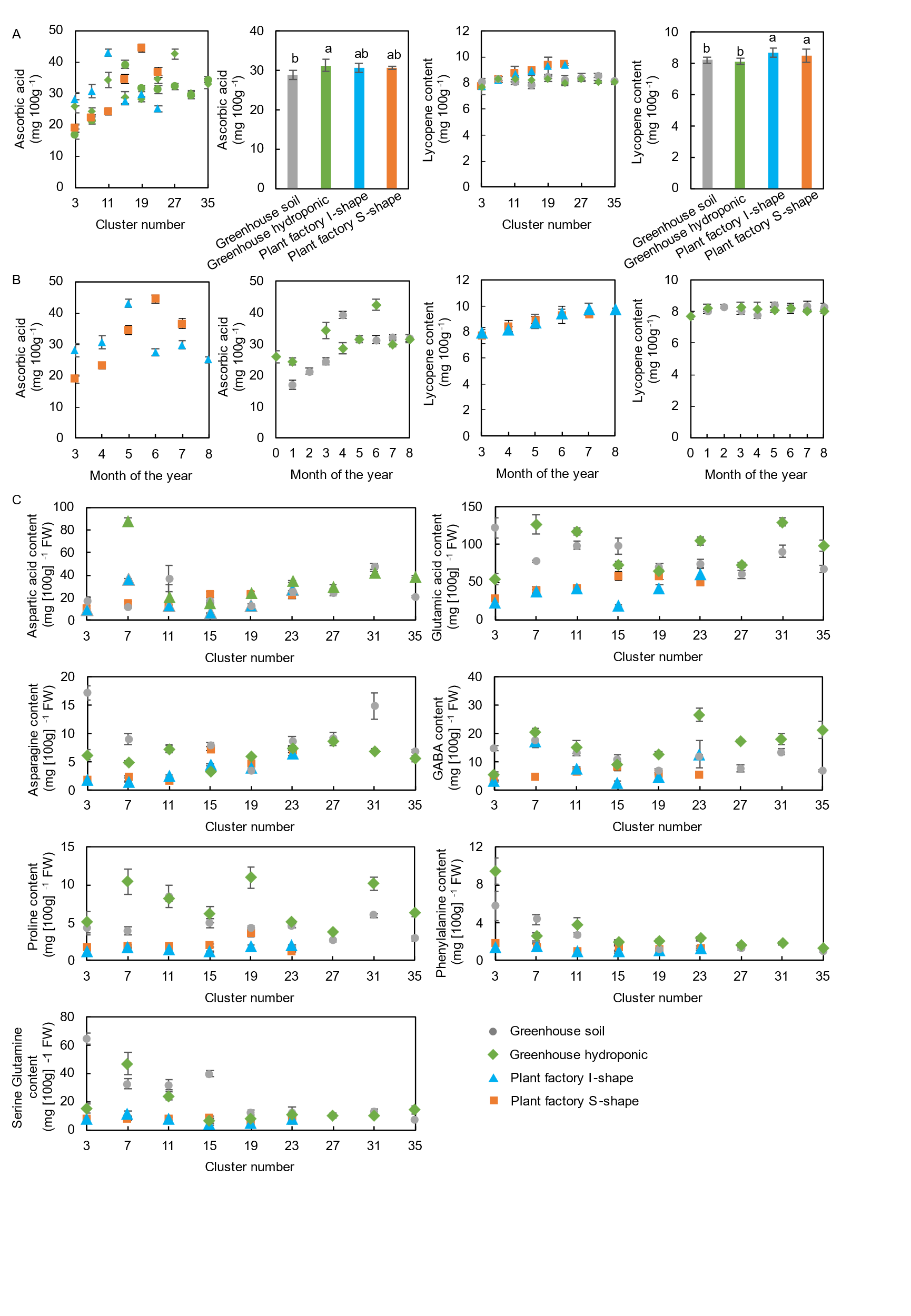

Figure S5

Functional nutrients of tomato from different flower cluster and different time of a year. A. Ascorbic acid and lycopene contents of tomato from different cluster and average contents. B. Ascorbic acid and lycopene contents of tomato at different time of a year and average contents. C. Amino acid contents of tomato from different cluster of. Data is mean ± SE. Bars with the same letter are not significantly different n = 50-90.

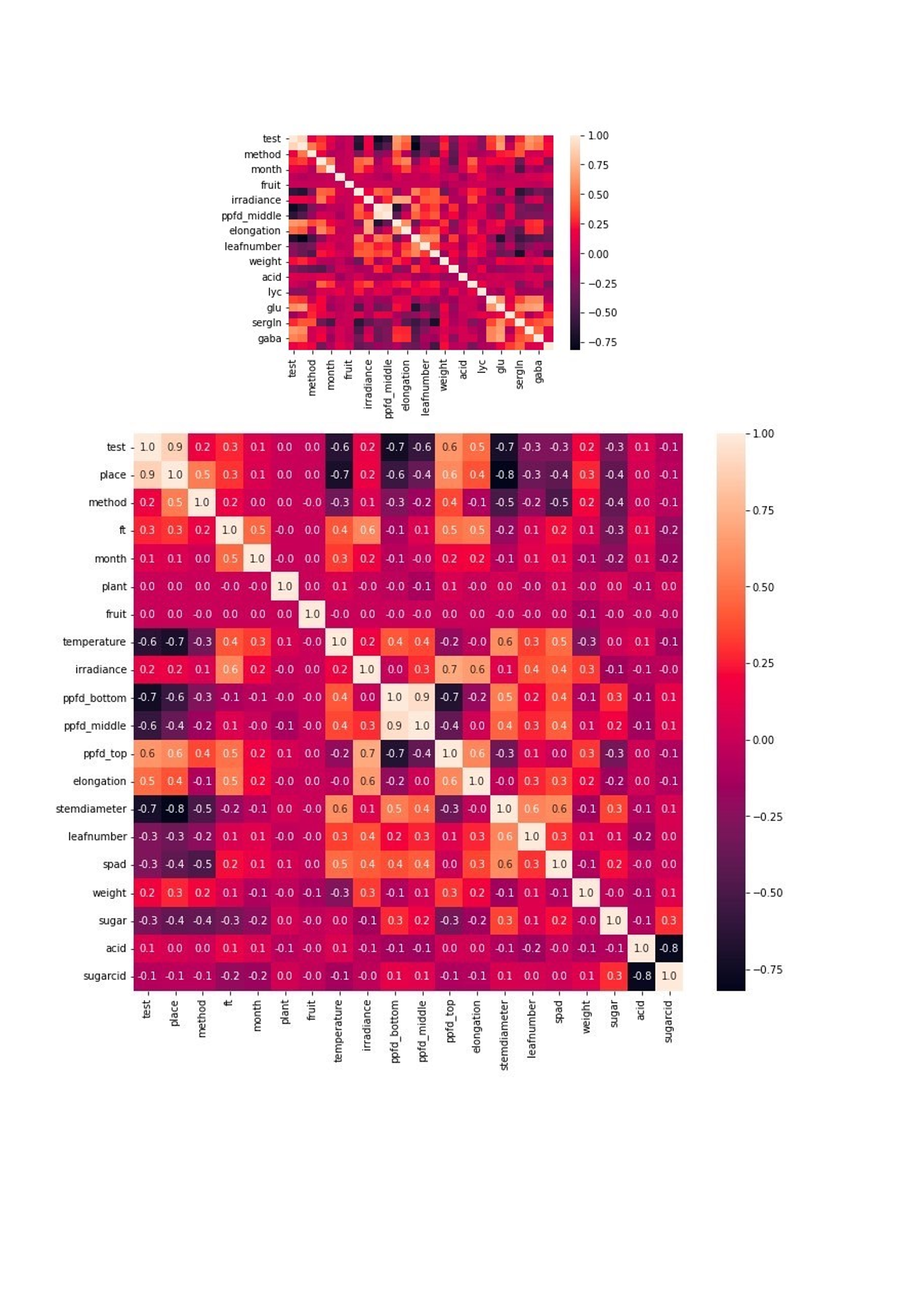

Figure S6

Correlation of different parameters in tomato plants and fruits cultivated/yielded in different conditions. Color represented correlation efficiency from 0.75 (darker color) to 1.0 (brighter color).

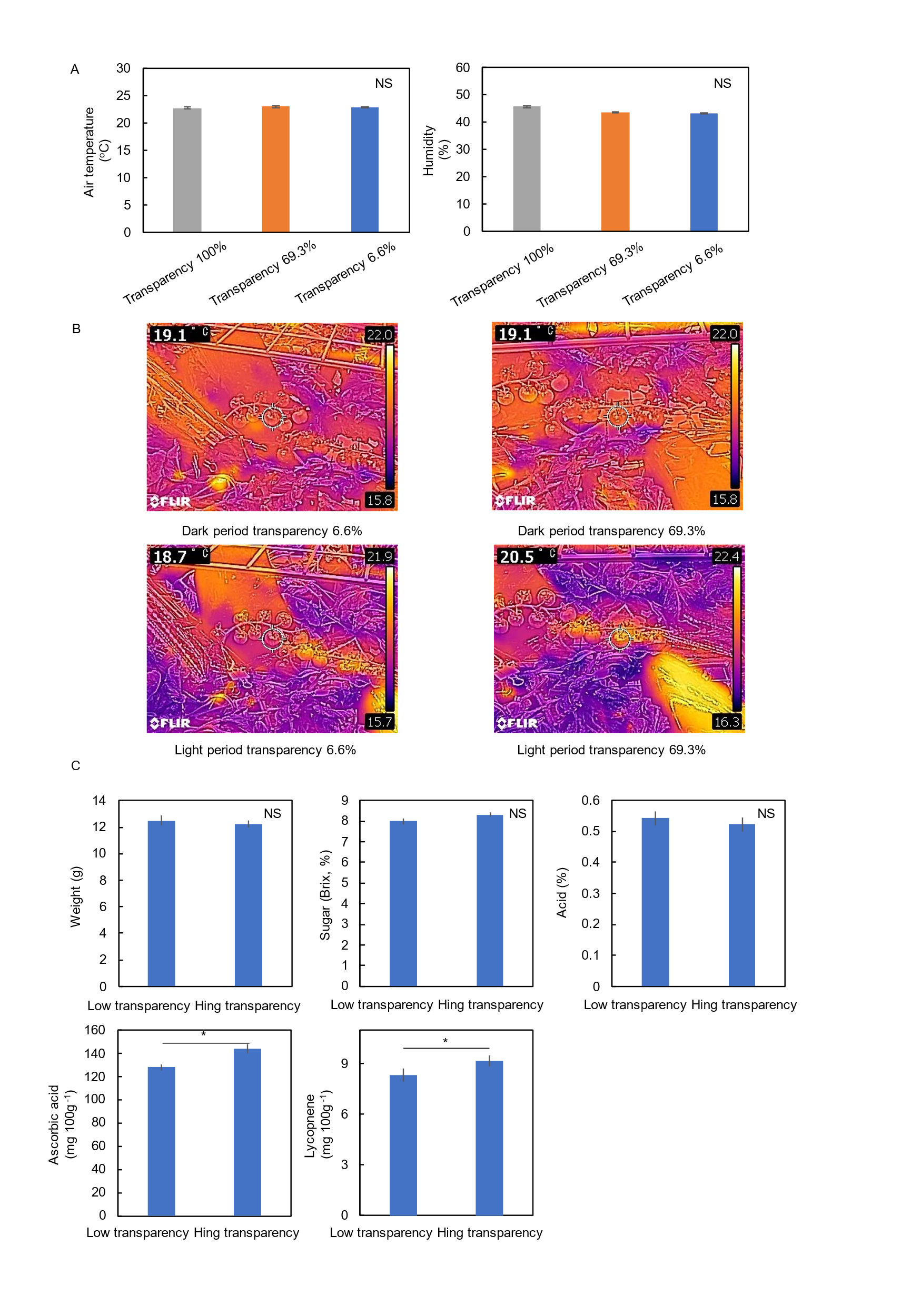

Figure S7

Environmental and yield data of tomato fruit clusters under the cover of membranes with different transparency. A. Air temperature and environmental humidity around tomato plants covered with membranes of different transparency. B. Thermographic photos of fruit clusters covered by membranes of different transparency. C. Yield parameters of tomato fruits from clusters covered by membranes of different transparency. Data is mean ± SE. Bars with the same letter are not significantly different n = 8.

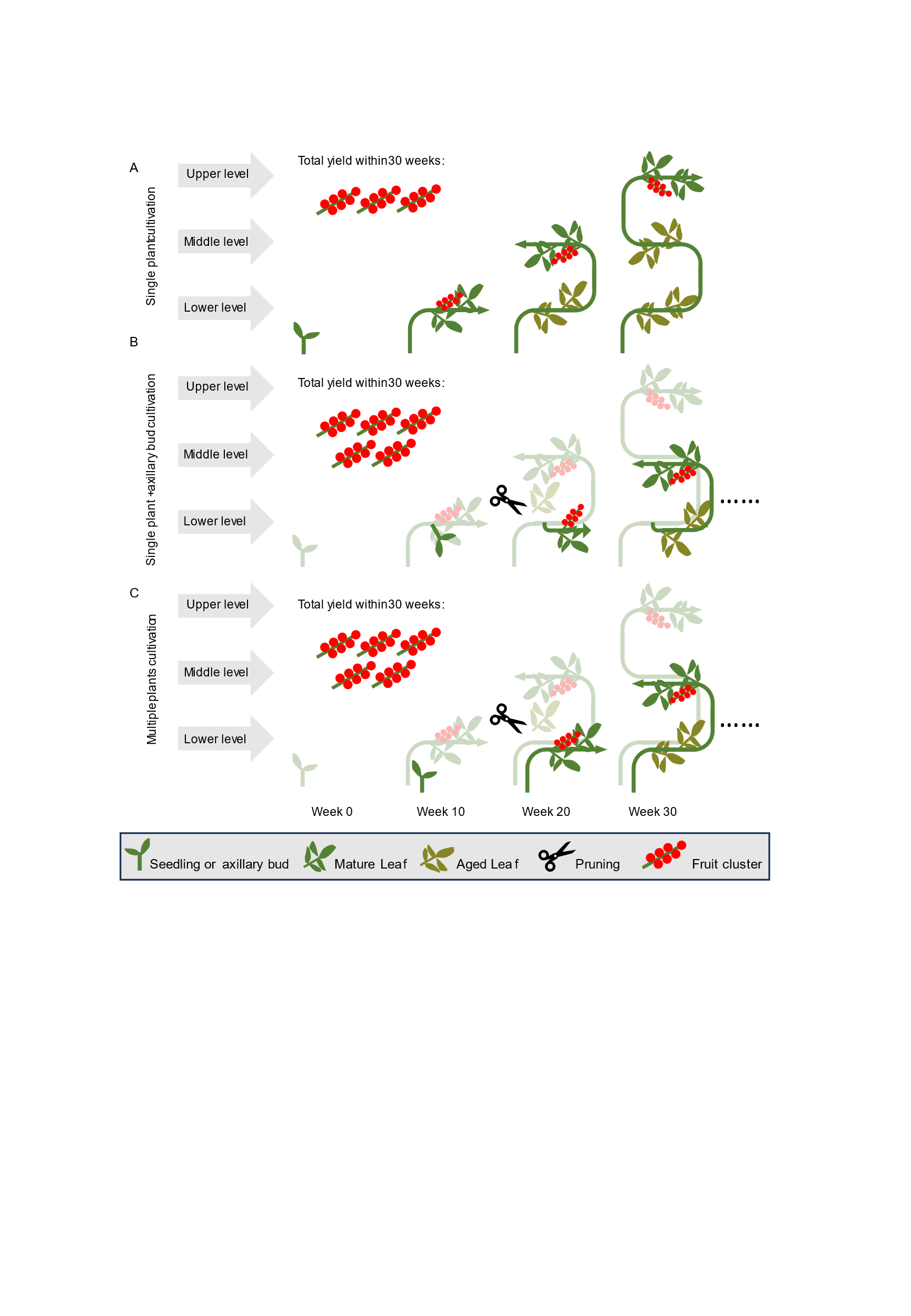

Figure S8

Use axillary buds or multiple plants cultivation to optimize production in a same S-shape cultivation system. A. Cultivation system using only 1 plant. B. Cultivation system using axillary bud at lower location to introduce a secondary shoot apex. C. Cultivation system using second plant.
